# Lack of correlation between number of neurons and behavioral performance in Swiss mice

**DOI:** 10.1101/428607

**Authors:** Neves Kleber, Gerson Duarte Guercio, Anjos-Travassos Yuri, Costa Stella, Perozzo Ananda, Montezuma Karine, Herculano-Houzel Suzana, Panizzutti Rogério

## Abstract

Neuronal number varies by several orders of magnitude across species, and has been proposed to predict cognitive capability across species. Remarkably, numbers of neurons vary across individual mice by a factor of 2 or more. We directly addressed the question of whether there is a relationship between performance in behavioral tests and the number of neurons in functionally relevant structures in the mouse brain. Naïve Swiss mice went through a battery of behavioral tasks designed to measure cognitive, motor and olfactory skills. We estimated the number of neurons in different brain regions (cerebral cortex, hippocampus, olfactory bulb, cerebellum and remaining areas) and crossed the two datasets to test the a priori hypothesis of correlation between cognitive abilities and numbers of neurons. As previous evidence indicates that environmental enrichment may increase neurogenesis and improve neuronal survival, we added a control group that did not undergo cognitive testing to rule out the possibility that our test battery could alter the neuronal number. We found that behavioral testing did not change numbers of neurons in the cerebral cortex and in the hippocampus. Surprisingly, performance in the behavioral tasks did not correlate strongly with number of neurons in any of the brain regions studied. Our results show that whereas neuronal number is a good predictor of cognitive skills across species, it is not a predictor of cognitive, sensory or motor ability across individuals within a species, which suggests that other factors are more relevant for explaining cognitive differences between individuals of the same species.

## Introduction

Brain size varies by more than 100,000-fold across species (Haug, 1987), and it has long been expected that this variation is related to the animal’s cognitive skills and behavioral flexibility. For example, behavioral innovation rate, a measure derived from a systematic collection of field notes of previously unreported behaviors, shows a positive correlation with forebrain size across bird species (Overington et al., 2009). Deaner and colleagues (2007) found that within the primate order, absolute brain size is a good predictor of a global cognition index extracted from meta-analyses. In a large, multi-group, coordinated effort to study many different species (from elephants to birds), MacLean and colleagues (2014) showed that absolute brain size is the best neuroanatomical predictor of performance in a task of self-control, the ability to inhibit a prepotent but ultimately counterproductive behavior. Across carnivoran species, problem-solving ability is also correlated with brain size (Benson-Amram et al., 2016).

There is evidence that brain size may also be an indicator of cognitive ability across individuals within a same species. For instance, it has been reported that in rats, a measure of a general intelligence factor (g) correlates positively with brain size (Anderson, 1993). Even within humans, a large meta-analysis showed that brain size correlates with IQ, albeit explaining only ~6% of the variance (McDaniel, 2005). Another study found that brain volume explains ~34% of the variance in general verbal ability in women (Witelson et al., 2006). A cross-sectional study showed that cerebellum volume explains some of the variance in g, even after controlling for frontal lobe volume (Hogan et al., 2011). And if we do not restrict ourselves to measures of cognition, still, in general, there is evidence that the size and cellular composition of functionally relevant structures are related to behavioral performance. One study found evidence that the size and number of cells in the frontal lobe of humans decrease with aging, mirroring motor function deterioration (Andersen et al., 2003). In a different context, one study found that the number of neurons in two song-related regions of the zebra finch brain correlates with their song repertoire size (Ward et al., 1998).

Brain size was once considered a proxy for the number of brain neurons both across and within species, based on assumptions about universal scaling rules of neuronal density and uniform surface densities of neurons within and across cortical areas (Rockel et al., 1980; Carlo and Stevens, 2013). More neurons, in turn, would determine larger information processing capacity, learning and flexibility (Williams and Herrup, 1988; Dicke and Roth, 2016). Recent evidence suggests that brain mass and number of neurons do not scale in the same way across species (reviewed in (Herculano-Houzel et al., 2014)) and that within a species, they are not correlated at all (Herculano-Houzel et al., 2015b). In light of these findings, it has been proposed that numbers of neurons in the cerebral cortex - and not brain size as a proxy for them - are the main neuroanatomical determinant of cognitive skills and flexibility across species (Harrigan and Commons, 2014; Dicke and Roth, 2016; Herculano-Houzel, 2017). The question then arises as to whether the same relationship holds true within a species.

Similarly aged individuals of a non-isogenic mouse strain exhibit variation by a factor of 1.33-fold in brain size and 1.63-fold in number of brain neurons (ratio between maximum and minimum values; (Herculano-Houzel et al., 2015b)). Until recently (Herculano-Houzel and Lent, 2005), it was impractical to estimate numbers of neurons in large numbers of individuals, and few studies addressed directly the relationship between an animal’s cognitive performance and its number of neurons. The ones that did address it made use of induced changes in the number of neurons, such as maternal treatments with growth hormone (Block and Essman, 1965; Zamenhof et al., 1966) and induced tetraploidy in salamander embryos (Vernon et al., 1955; Vernon and Butsch, 1957). Although such studies showed that manipulations resulting in altered number of neurons also changed cognitive performance, they did not address whether numbers of neurons are a predictor of cognitive or motor ability in healthy and normally developed animals.

Here, we sought to address this question directly: is there a relationship between naïve (untrained) performance in behavioral tests and the number of neurons in functionally relevant structures of the brain? To this end, we took Swiss mice through a series of behavioral tasks designed to measure cognitive, motor and olfactory skills and then we used the isotropic fractionator method to estimate the number of neurons in their brain regions. Specifically, we tested the following *a priori* hypotheses: that performance in an olfactory test and in the rotarod test correlates with number of neurons in the olfactory bulb and cerebellum, respectively; that performance in the Morris water maze (a navigation learning task) correlates with number of neurons in the hippocampus and cerebral cortex; that performance in operant training and in the puzzle-box (two tasks that require perceptual and executive functions) correlates with number of neurons in the cerebral cortex and hippocampus. In addition, as evidence indicates that environmental enrichment may increase neurogenesis and improve neuronal survival (Clemenson et al., 2015), we compared numbers of neurons in those animals to a control group of animals that did not undergo cognitive testing to rule out the possibility that the test battery could alter the brain cellularity.

## Methods

### Behavioral testing

All experiment procedures used in this study were approved by the Animal Care and Use Committee at Universidade Federal do Rio de Janeiro (protocol number 01200.001568/2013–87). □

Male mice were aged 2 months at the beginning of behavioral testing, and were naïve (inexperienced) in any type of behavioral tasks. All mice were housed in groups of 2 to 5 per cage. We performed the tests in the following order: olfactory test, rotarod, Morris water maze, puzzle-box and operant conditioning. To control for the effects of human manipulation and physical exercise, the control group underwent the physical part of each test, without the cognitive challenge, as explained below for each task. Mice were euthanized at the end of the behavioral testing when they were 4 months old.

### Olfactory Test

We used a modified version of the hidden peanut butter finding test (de Souza et al., 2011). On the first day, mice were food-deprived and given one peanut each to prevent bait shyness. On the next day, we exposed them to an arena of 40 × 40 cm filled with bedding, and a buried peanut in it. In three trials starting at different positions, we allowed mice 15 minutes to find the peanut. The latency to find the peanut in each of the three trials was recorded and summed, giving the total latency used as a score, so smaller scores indicate better performance. We exposed control animals to the same procedure but without the buried peanut in the arena.

### Rotarod

To assess balance and motor learning, we measured the latency to fall from a mouse rotarod (Insight Equipamentos, Brazil). On the first day, we exposed mice to a rotating rod starting at 14 RPM and accelerating until 35 RPM for a maximum of 4 minutes for trial. Mice underwent 5 trials and we recorded the latency to fall in three trials, after discarding the highest and lowest latencies. On the next day, mice underwent the same experiment for three trials, and the final score was the sum of the latency to fall over 6 trials in the first and second days. Control mice were exposed to the rotarod at a constant 14 RPM for a similar duration.

### Morris Water Maze

The maze (circular tub, 110 cm diameter) was filled with water at 22°C made opaque by the addition of milk powder. All testing occurred under dim lighting. On the first day, we placed the animals on the hidden platform 3 times for 30 seconds each. On the second day, we added the extra-maze cues and gave the mice 6 opportunities to locate the platform (6 trials of 90 seconds maximum each). If a mouse did not find the platform during this time, we led it to the platform. In each trial, the mice stayed on the platform for 30 seconds, and the inter-trial interval was 10 min, in which we dried each animal briefly under a warm light before returning them to their cages. On the third day, each mouse had four more opportunities to find the platform. After 90 minutes of the last trial, we returned each mouse to the maze for 90 seconds, this time without the platform. Anymaze Video Tracking Software was used to record swimming patterns, and we divided the maze into four quadrants. The “goal” quadrant was the one in which the platform was located. The score was calculated by subtracting the amount of time a mouse spent on the goal quadrant from the time it spent in the opposite quadrant. We lost the data from 6 mice because of a technical problem with the video camera. We allowed control mice to swim in similar intervals without the cues and the platform.

### Puzzle-box

The test was adapted from (Ben Abdallah et al., 2011). Briefly, the arena consisted of a wooden white box divided by a removable barrier into two compartments: a brightly-lit start zone (58 cm long, 28 cm wide) and a smaller covered goal zone with bedding and food (15 cm long, 28 cm wide). Lightly food-deprived mice always started in the brightly-lit zone, and were given a 5 minute opportunity to go to the goal zone three times per day, undergoing a total of nine trials (T1–T9) over 3 consecutive days. Over the nine trials, they were challenged with obstructions of increasing difficulty placed at the underpass. On day 1, the underpass was unblocked in T1, and the barrier had an open door over the location of the underpass. In T2 and T3 the barrier had no doorway and mice entered via the small underpass. On day 2, T4 was identical to T2 and T3, but on T5 and T6, however, the underpass was filled with sawdust and mice had to dig their way through (burrowing puzzle). On day 3, T7 was a repetition of T5 and T6, and on T8 and T9, mice were presented with the plug puzzle, with the underpass obstructed by a cardboard plug that mice had to pull with teeth and paws to enter the goal zone. This sequence allowed assessing problem-solving ability (T5 and T8), learning/short-term memory (T3, T6, and T9), while the repetition on the next day provided a measure of long-term memory (T4 and T7). The total latency was used as a readout of the puzzle-box test, so smaller scores indicate better performance. Data from one mouse was excluded because it did not reach the goal zone in T1, which indicated lack of motivation to complete the task. Control mice were exposed to the arena and lightly food-deprived as in T1.

### Operant Conditioning

For 32 days, testing consisted of different phases that involved the following cognitive capacities: auditory tone discrimination, attention, memory, problem solving and cognitive flexibility. All testing sessions consisted of 1 hour per day, except the second phase, which consisted of two sessions of 30 minutes per day. To motivate mice to participate, they were lightly food-deprived in order to maintain around 90% of the baseline weight. In the first phase of testing, they got a food reward for making a “go” response. The reward consisted of a 20 mg food pellet (BioServe, Frenchtown, NJ). After making more than 40 responses for two sequential days, the mice went to the second phase, in which they had to make a go response only during a 3 s window after the presentation of a 600 ms tone. Mice graduated after making a correct go response at least 70% of the time again for two sequential days, which was the same criteria for the following phases. On the next phase, there were 3 different sounds and mice had to figure out the correct one. In the fourth phase, mice had to find a correct tone in a new set of 3 different sounds of 200 ms each. Finally, the fifth phase consisted of a 1-back, in which mice had to perform a go response when a 200 ms tone A was played following another specific 200 ms tone B. All tones were presented at 70 dB SPL. Testing was performed in an acoustically transparent operant training chamber contained within a sound-attenuated chamber. The performance was calculated multiplying the number of days spent on each phase by its number and summing the results (if an animal stayed 16 days in phase 1 and 16 days in phase 2, the score would be (16*1) + (16*2) = 48). Therefore, mice that reached higher phases faster over the 32 days of testing got higher scores.

### Perfusion

We analyzed the cellular composition of the brain of 32 mice that performed the behavioral testing (in four batches spaced a few months apart), and 27 mice in the control condition. Animals were 4 months old at the time of death. Animals were killed by an overdose of ketamine through an IP injection and perfused through the heart with a 0.9% saline solution followed by 4% phosphate-buffered paraformaldehyde (PFA). All animals from the same batches were killed and perfused on the same day.

### Dissection

The brains were removed from the skull, cleaned of dura-mater and major vasculature and then left immersed in PFA for exactly two weeks of post-fixation. After this, they were dissected into cerebral cortex, hippocampus, cerebellum, olfactory bulb and remaining areas (brainstem, diencephalon and basal ganglia). The cerebral cortex was further divided into anterior and posterior regions. The anterior cortex was defined as the regions anterior to the genu of the corpus callosum on the anterior-posterior axis. All structures were weighed immediately after dissection. The left and right halves of the cerebral cortex (anterior and posterior) and hippocampus were processed separately, while for the cerebellum, remaining areas and olfactory bulb, both halves were processed together.

### Isotropic Fractionator

After post-fixation, brains were processed to obtain estimates on their number of neuronal and non-neuronal cells, using the isotropic fractionator (Herculano-Houzel and Lent, 2005). The method consists of using mechanical dissociation to transform heterogeneous brain tissue into a suspension of cell nuclei that can be kept homogeneous by agitation. Nuclei can then be counted in samples from the suspension and stained by immunocytochemistry. The isotropic fractionator has been shown by two independent groups to give estimates comparable to stereological estimates (Bahney and von Bartheld, 2014; Miller et al., 2014; Herculano-Houzel et al., 2015a).

Structures were mechanically dissociated in Triton X-100 1% in a 40 mM solution of sodium citrate. The resulting suspension with all nuclei was stained with the fluorescent DNA marker DAPI (4’-6-diamidino-2-phenylindole dihydrochloride, Invitrogen, USA) diluted 1:20 from a stock solution of 20 mg/L and brought up to a readable volume in a graduated tube with phosphate-buffered saline (PBS). The density of nuclei in the suspension was estimated by counting at least 4 samples of the suspension in a Neubauer counting chamber under a fluorescence microscope. The coefficient of variation between samples was typically below 0.10. Once the estimates for cell number were obtained, a small sample of 1 mL was taken from the suspension to undergo immunocytochemistry for a neuronal nuclear antigen (NeuN, Millipore mab377), expressed in the nuclei of most neuronal cell types and not in non-neuronal cells (Mullen et al., 1992; Gittins and Harrison, 2004). This allowed us to estimate the absolute number of neuronal cells in the tissue. The number of non-neuronal cells was determined by subtraction. All counts were made in the same day or the next after the immunocytochemistry, to preserve the fluorescence and avoid one source of variability in the estimates.

### Statistics

All statistical analyses were performed in R 3.4.1 (R Core Team, 2017). Performance in the behavioral tasks was ranked and the ranks were considered in the analyses. Pairwise correlations between the cellular composition of brain structures and the performances in the behavioral tasks were calculated using Spearman rank correlation, allowing us not to miss non-linear associations between the variables. We chose an alpha level of 5% for statistical significance.

## Results

First, we investigated whether performance between tasks was correlated. We hypothesized a priori that the three tasks that depend the most on higher cognitive functions (puzzle-box, operant conditioning and Morris water maze) would have positively correlated performances. However, all correlations were non-significant, except for Hidden Peanut Butter Test x Puzzle Box **(Table 1)**. Therefore, in the analysis, we used the individual scores.

**Table 1.** Correlations between individual performance across different tasks. Each cell in the table shows the Spearman correlation and 95% confidence interval between behavioral performance of the same animal on two different tasks. No correlations but one (Hidden Peanut Butter Test x Puzzle Box) were significant at the 5% level. Sample size is 26–32, depending on the comparison.

|  | Hidden Peanut<br>Butter Test | Morris Water<br>Maze | Operant<br>Conditioning<br>AUC | Puzzle Box | Rotarod |
| --- | --- | --- | --- | --- | --- |
| Rotarod | 0.23<br>[-0.13, 0.54] | -0.34<br>[-0.64, 0.05] | 0.13<br>[-0.23, 0.46] | -0.03<br>[-0.37, 0.32] | 1 |
| Puzzle Box | 0.50<br>[0.18, 0.73] | 0.02<br>[-0.37, 0.40] | 0.14<br>[-0.23, 0.47] | 1 | -0.03<br>[-0.37, 0.32] |
| Operant Conditioning<br>AUC | -0.11<br>[-0.45, 0.25] | 0.17<br>[-0.24, 0.52] | 1 | 0.14<br>[-0.23, 0.47] | 0.13<br>[-0.23, 0.46] |
| **Morris Water Maze** | -0.10<br>[-0.47, 0.30] | 1 | 0.17<br>[-0.24, 0.52] | 0.02<br>[-0.37, 0.40] | -0.34<br>[-0.64, 0.05] |
| **Hidden Peanut Butter Test** | 1 | -0.10<br>[-0.47, 0.30] | -0.11<br>[-0.45, 0.26] | -0.50<br>[0.18, 0.73] | 0.23<br>[-0.13, 0.54] |

We then looked at the variation in our measures: behavioral performance and neuronal numbers, to guarantee that any lack of correlation is not an artifact of low variance in our sample. In **(Figure 1)** we show the variation in the behavioral performances. They all vary by more than one standard deviation around the mean. For neuronal numbers, besides investigating the variation in the sample, we also compared our results with a previous study of our group that measured the intraspecific variation in neuronal numbers, in the same lineage of mice (Swiss; Herculano-Houzel et al., 2015). As shown in **(Figure 2)**, the mass of the mice brain structures in these two studies largely overlap. The larger amount of variation in the olfactory bulb in both studies is attributed to variation in dissection and weighing of this very small structure. **(Figure 3)** shows that numbers of neurons are also similar between the two studies. We performed F-tests to compare the variances between them. A significant difference was found only for the cerebellum (F = 0.310, p < 0.001) and rest of brain (F = 0.215, p < 0.001), where the variance in our study was smaller than that of Herculano-Houzel and colleagues (2015b). These results indicate that there are no systematic differences in both studies.

**Figure 1:**
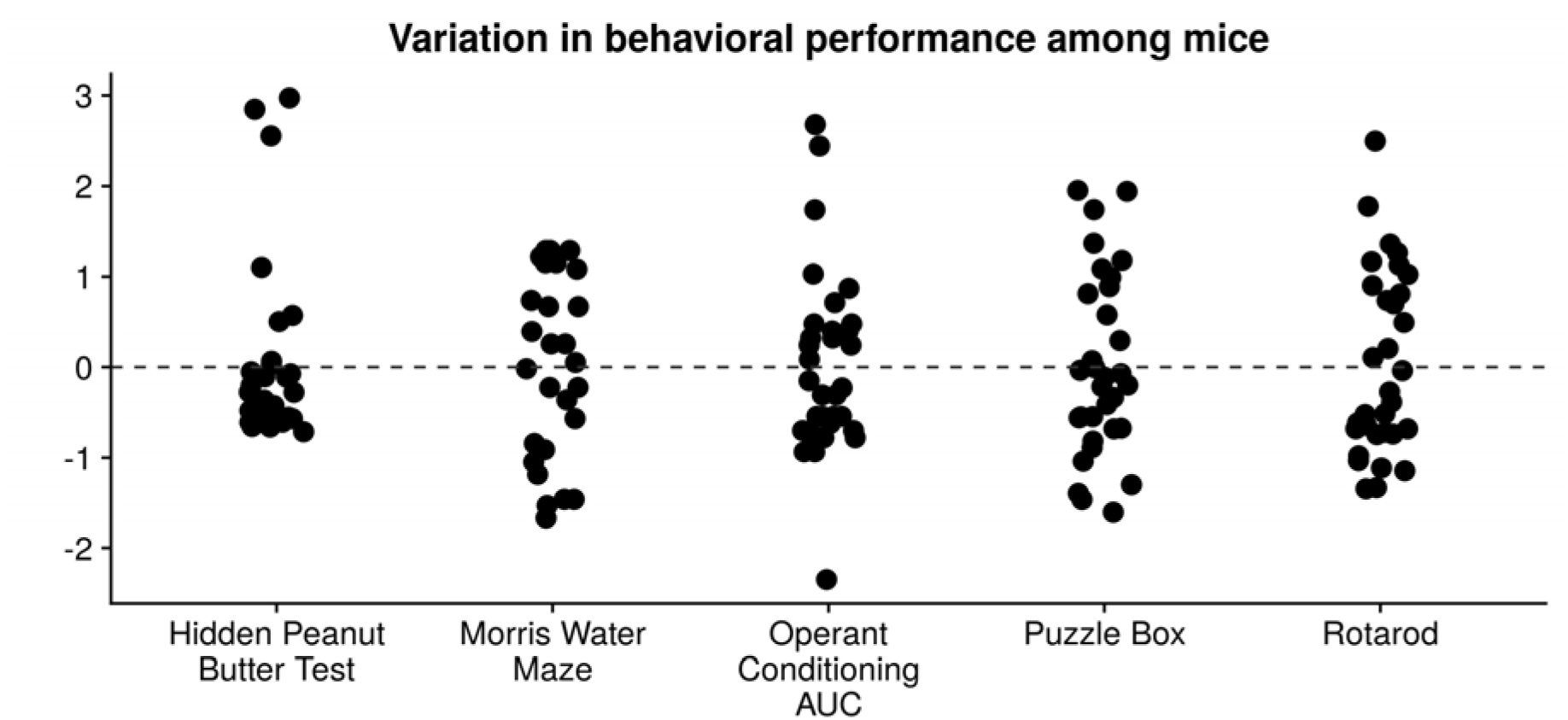
Variation in behavioral performance for each task, within our sample. Shown are the scores for each mouse in each task, standardized by subtracting the mean and dividing by the standard deviation. All of the scores show a variation of at least one standard deviation around the mean.

**Figure 2.**
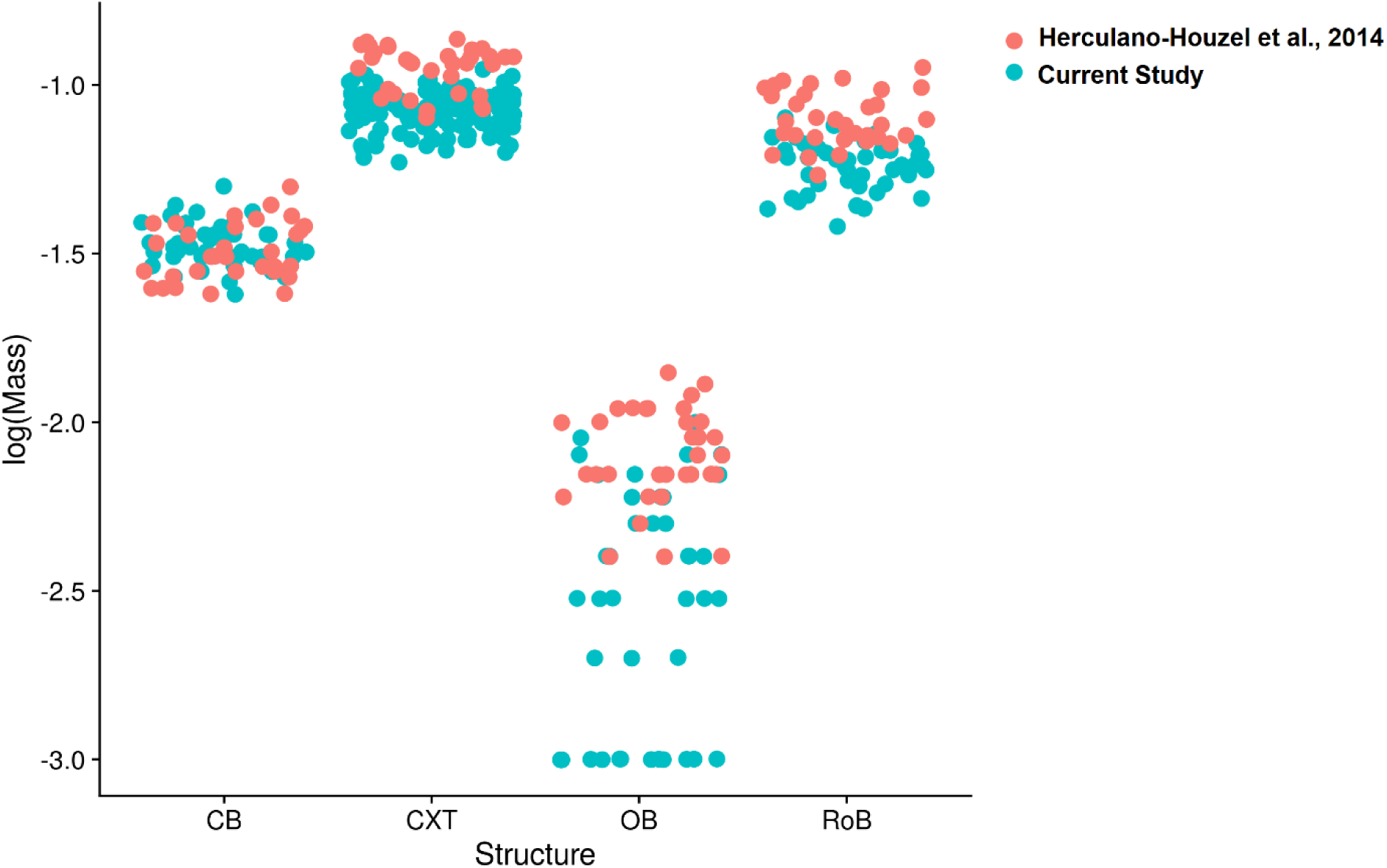
Comparison of structure mass in each structure in this and in a previous study. The range of the mass of the brain structures from the two studies overlap. CB, cerebellum; CXT, cerebral cortex; OB, olfactory bulb; RoB, rest of the brain. The larger amount of variation in the olfactory bulb in both studies is attributed to variation in dissection and weighing.

**Figure 3.**
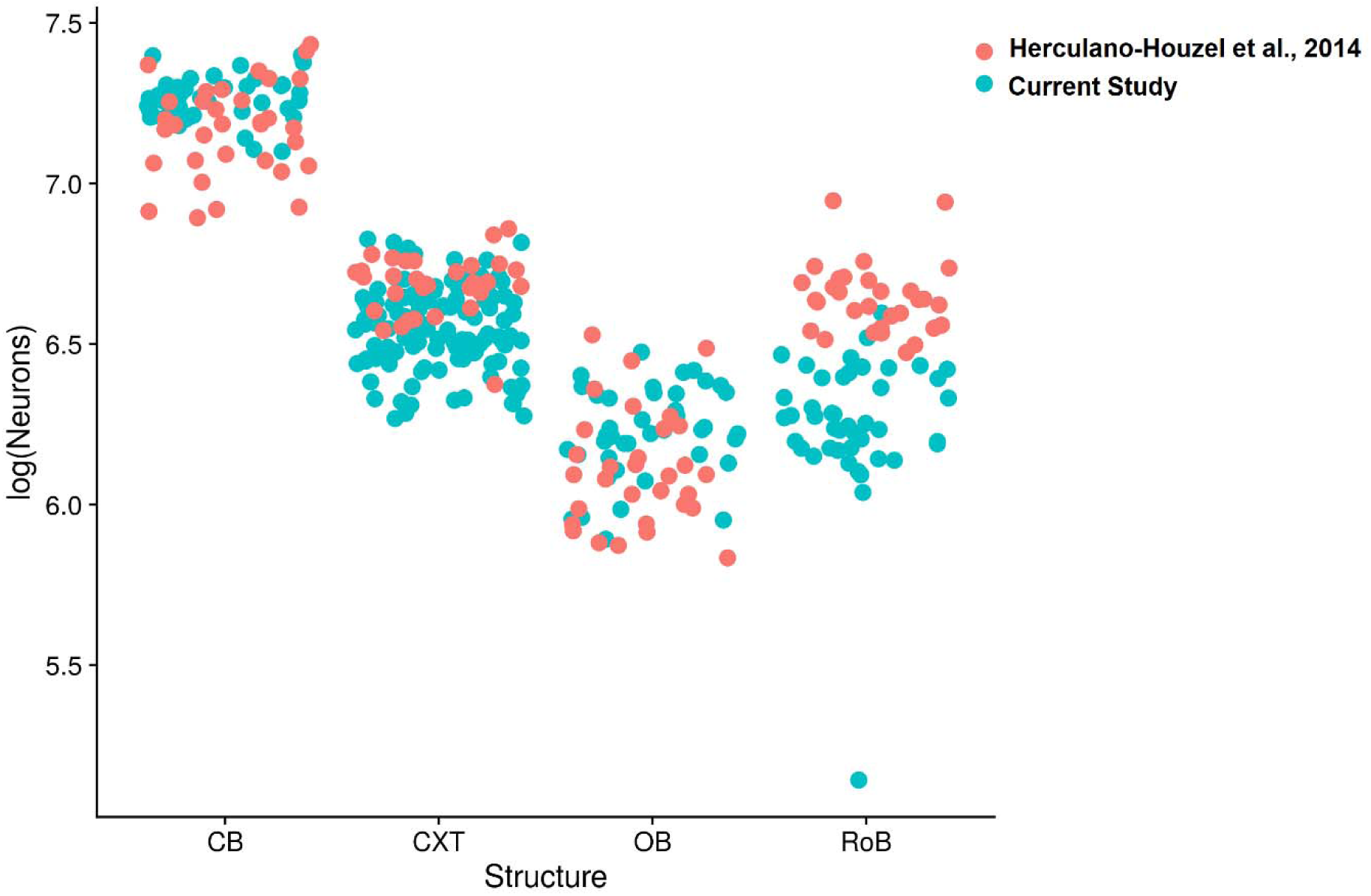
Comparison of number of neurons in each structure in this and in a previous study. Again, the range of number of neurons in brain structures from the two studies mostly overlap. CB, cerebellum; CXT, cerebral cortex; OB, olfactory bulb; RoB, rest of the brain. A significant difference was found only for the cerebellum (F = 0.31, p < 0.001; smaller in the current dataset) and rest of brain (F = 0.25; p < 0.001; larger in the current dataset).

The large variability across individual mice in performance in the various behavioral tests together with the large variability in numbers of neurons in the different brain structures allowed us to test our *a priori* hypotheses regarding the number of neurons and performance in functionally-relevant brain regions. We found no significant correlations between the number of neurons in the olfactory bulb and the latency to find the peanut in the olfactory test **(Figure 4)**, number of neurons in the cerebellum and latency to fall in the rotarod task **(Figure 5)**, number of neurons in the hippocampus and performance in the Morris Water Maze **(Figure 6)**, and number of neurons in the cerebral cortex and performance in the Morris Water Maze (**Figure 7)**, number of neurons in the hippocampus **(Figure 8)** and the cerebral cortex **(Figure 9)** with performance in the puzzle-box (**Figure 10)** and with performance in operant training **(Figure 11)**. Taken together, these results indicate that the number of neurons is not a predictor of individual performance in mice (see **Table 2** for all correlations). Similarly, we did not find any correlation between behavioral performance and numbers of non-neuronal cells in the relevant structures above **(Table 3)**.

**Figure 4.**
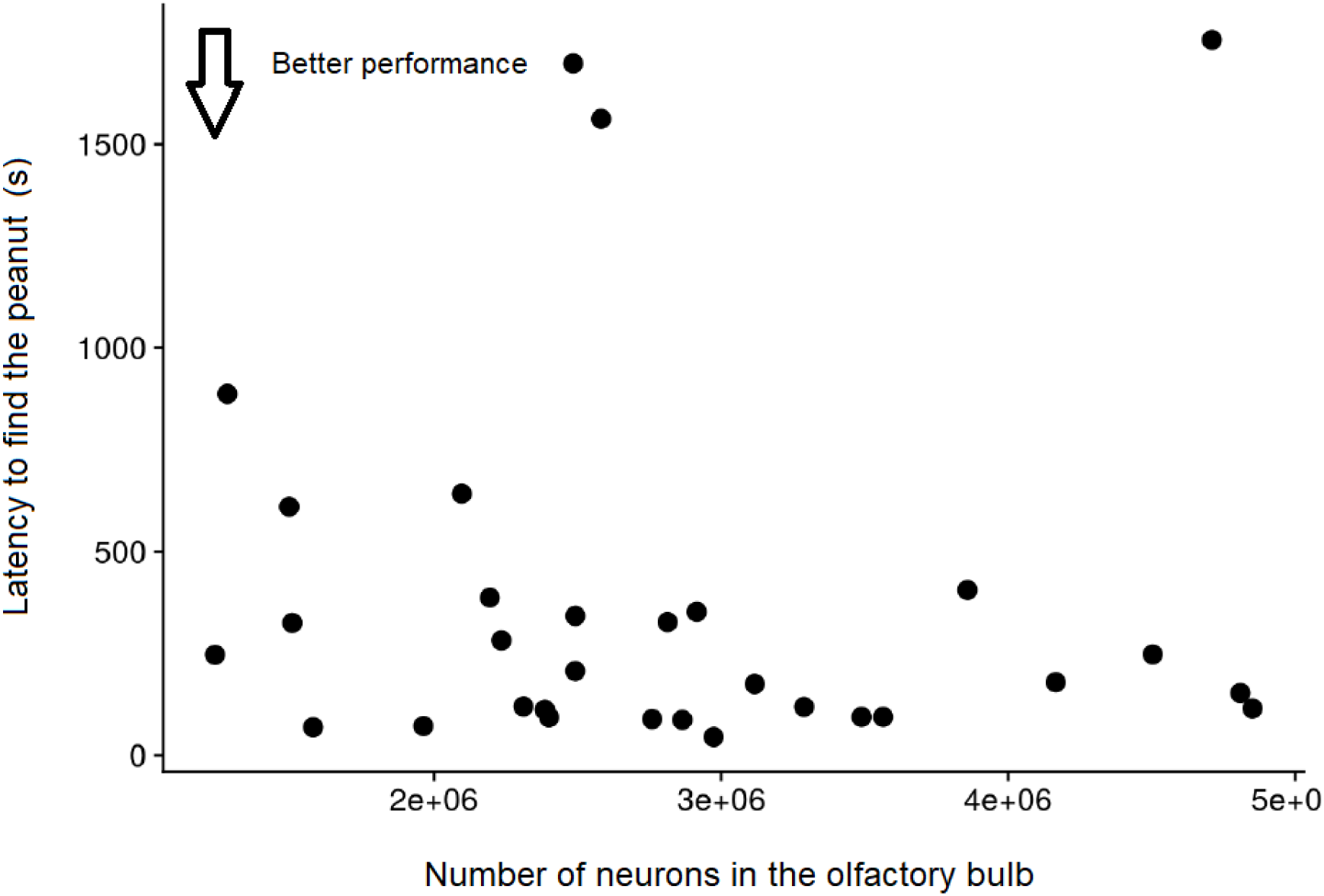
Lack of correlation between performance in the olfactory task and number of neurons in the olfactory bulb. (N = 30, ρ = −0.13, p = 0.98).

**Figure 5.**
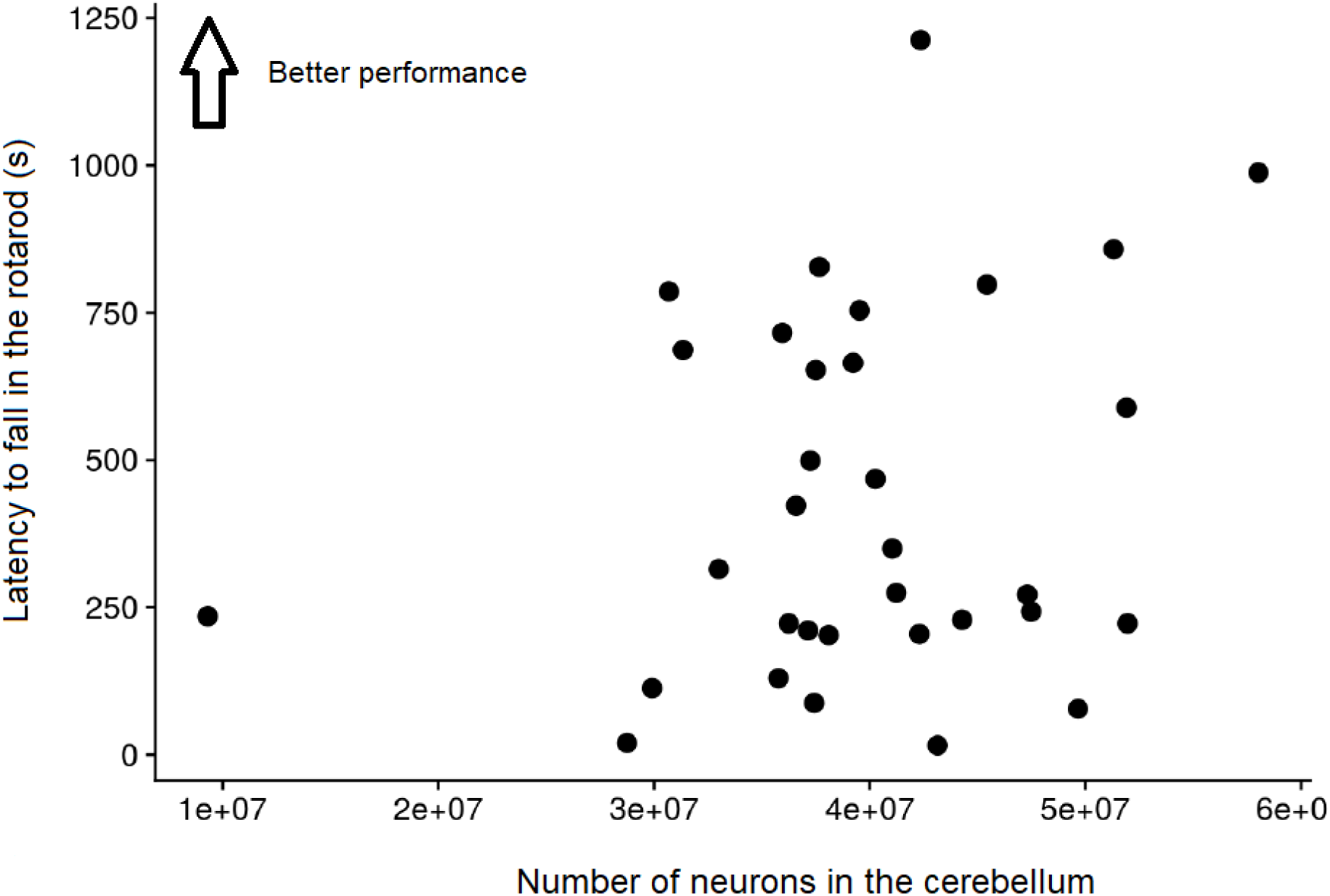
Lack of correlation between performance in the rotarod and number of neurons in the cerebellum. (N = 32, ρ = 0.18, p = 0.22).

**Figure 6.**
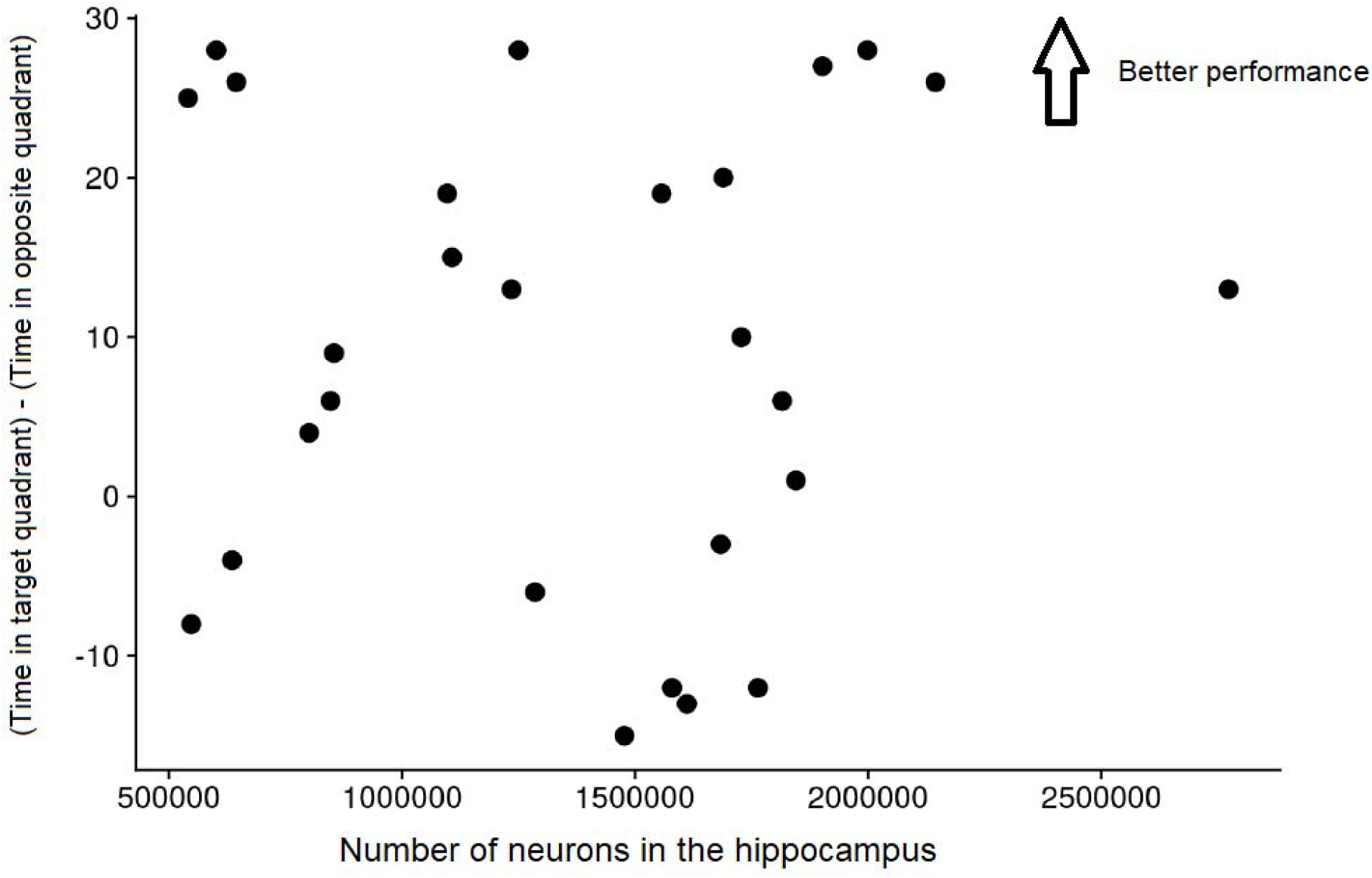
Lack of correlation between performance in the Morris water maze and number of neurons in the hippocampus. (N = 26, ρ = 0.03, p = 0.97)

**Figure 7.**
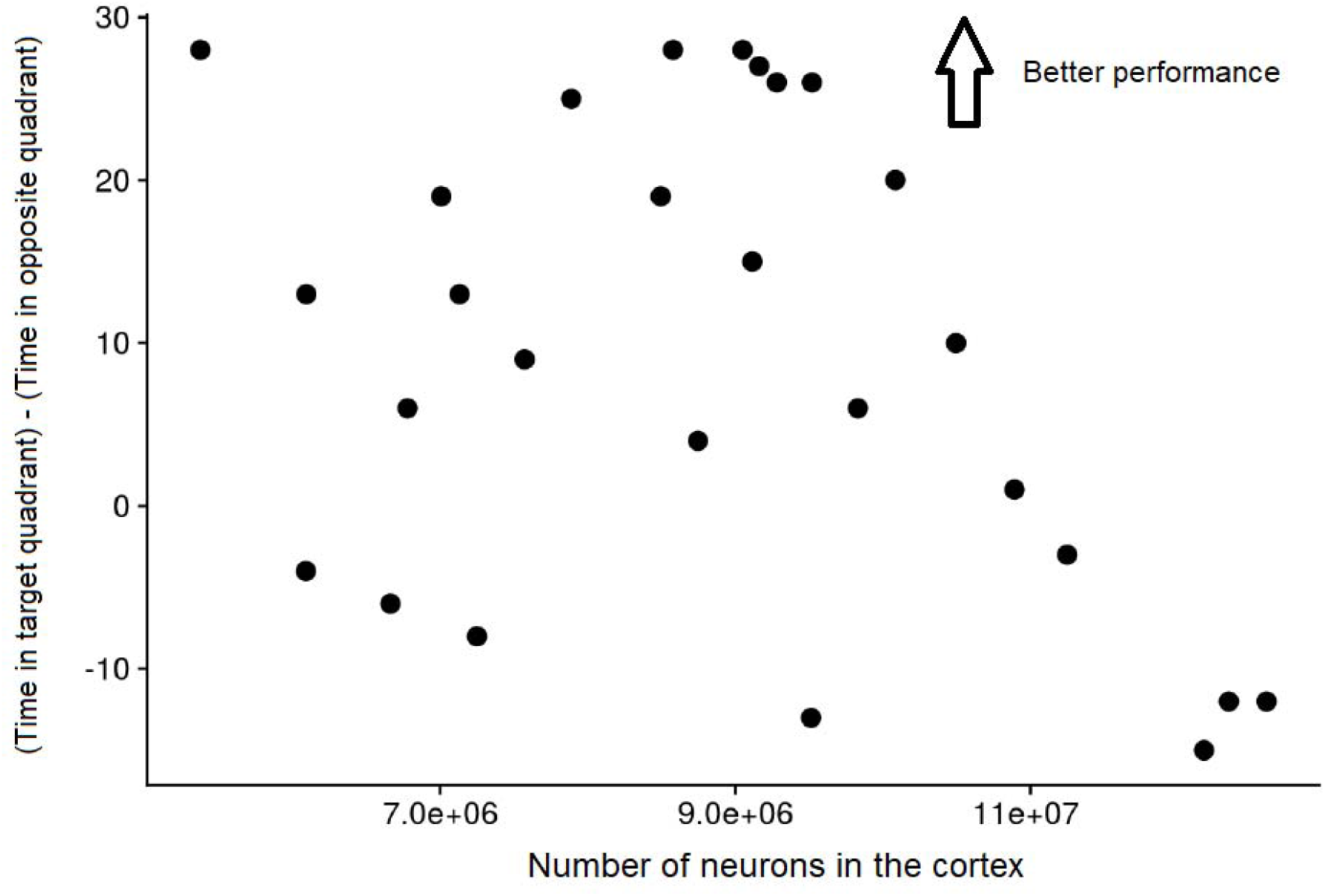
Lack of correlation between performance in the Morris water maze and number of neurons in the cerebral cortex. (N = 26, ρ = −0.29, p = 0.06).

**Figure 8.**
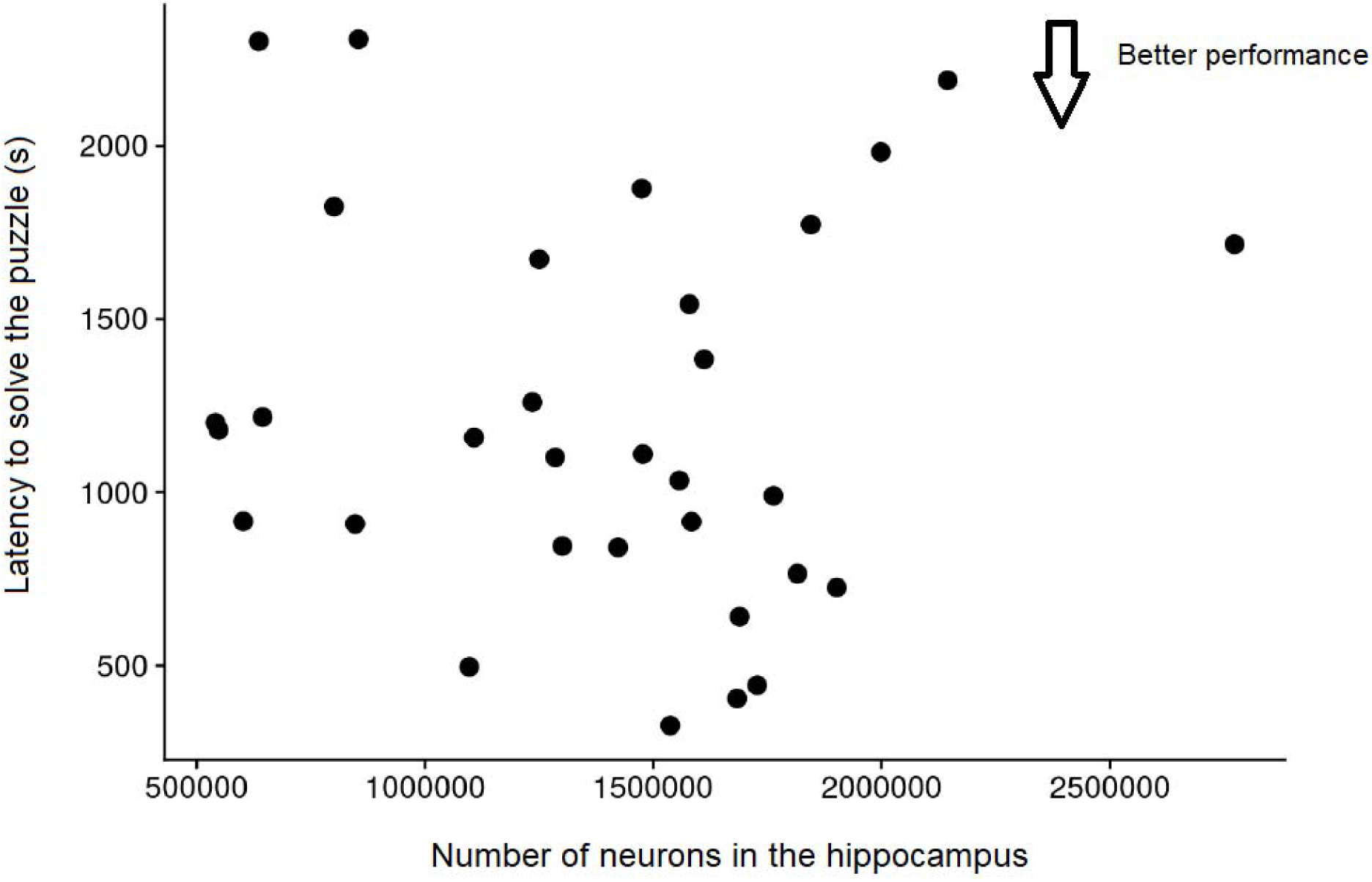
Lack of correlation between performance in the puzzle box and number of neurons in the hippocampus. (N = 31, ρ = −0.11, p = 0.87).

**Figure 9.**
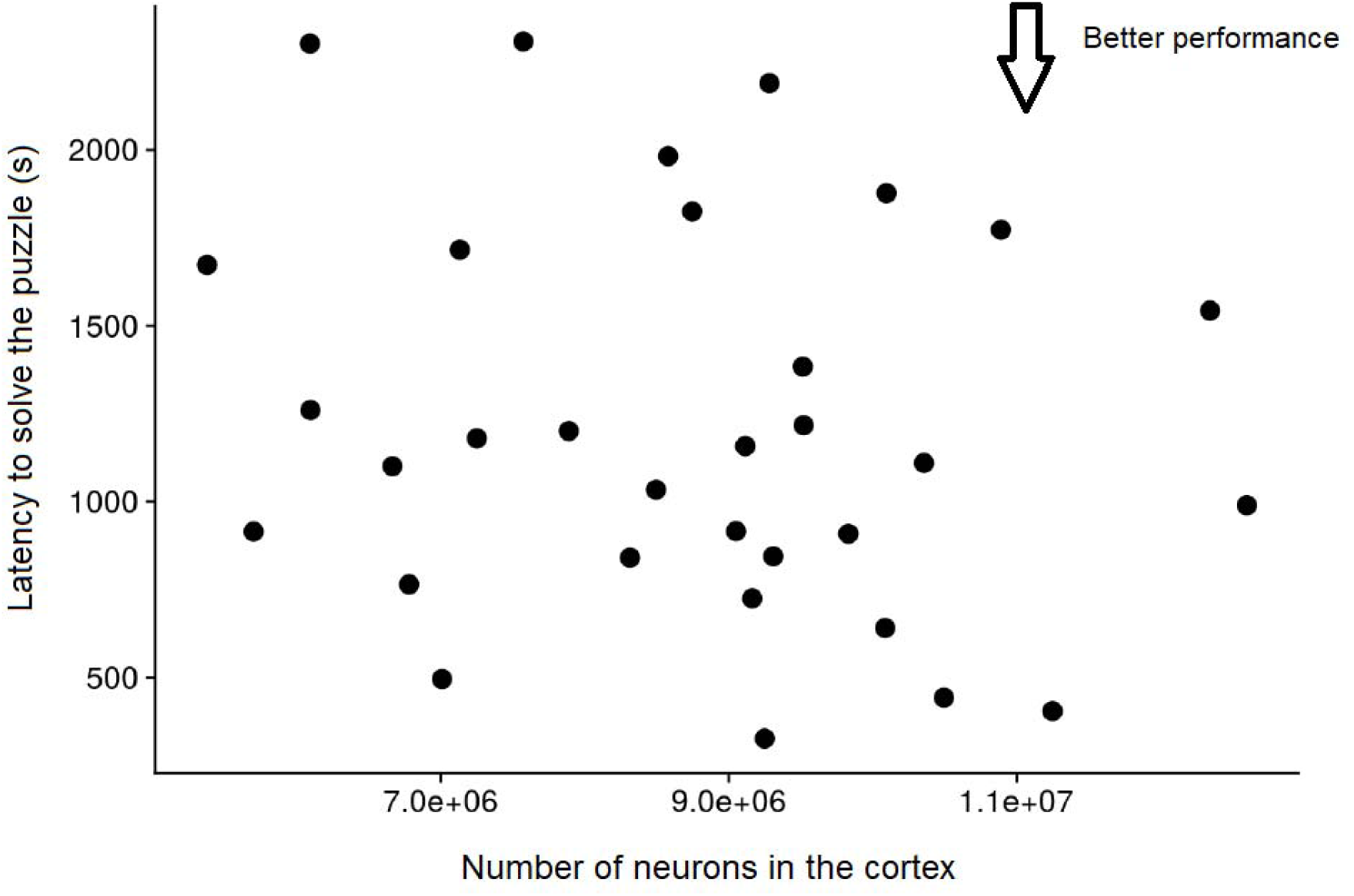
Lack of correlation between performance in the puzzle-box and number of neurons in the cerebral cortex. (N = 31, ρ = −0.17, p = 0.32).

**Figure 10.**
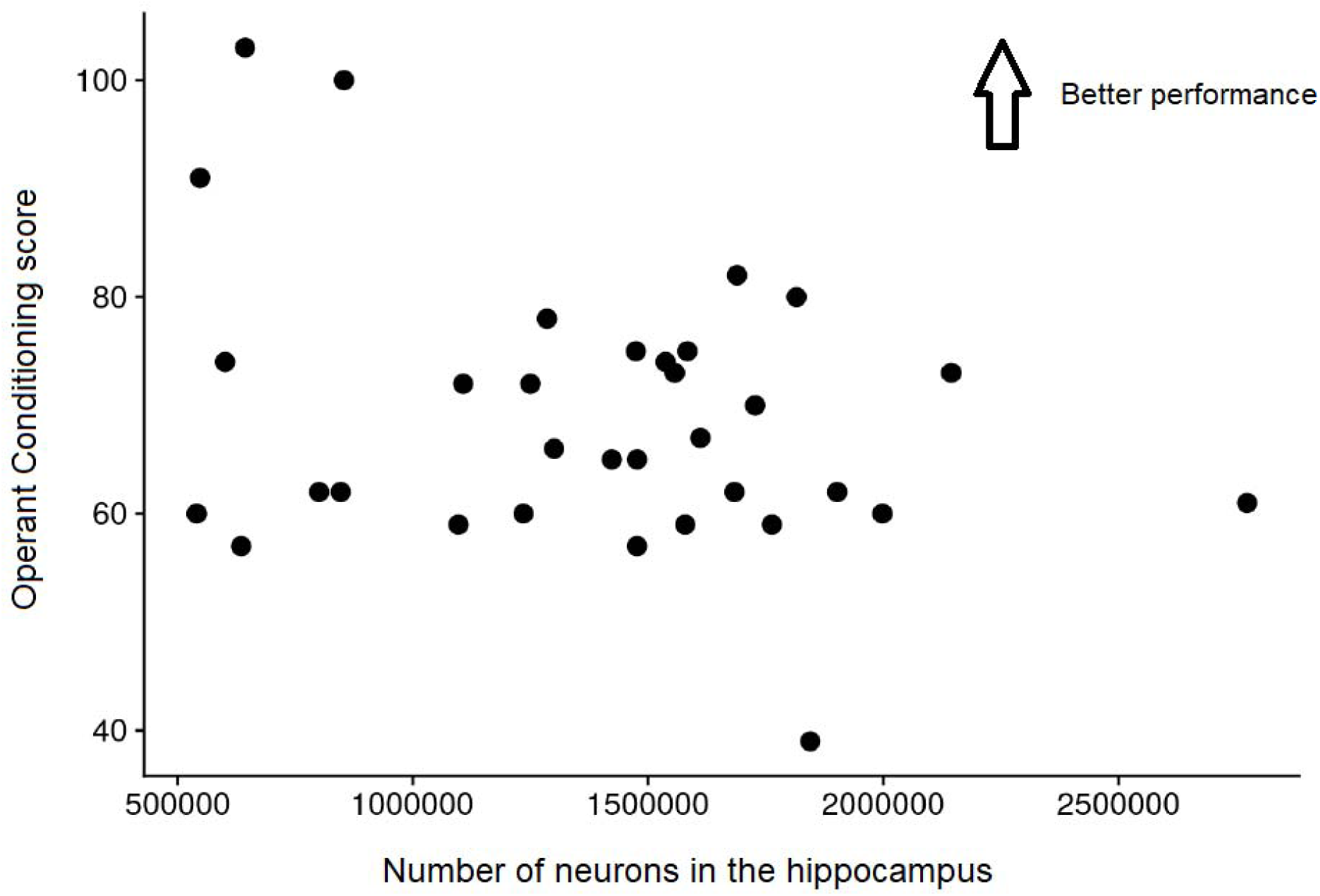
Lack of correlation between performance in the operant training and number of neurons in the hippocampus. (N = 32, ρ = −0.14, p = 0.07).

**Figure 11.**
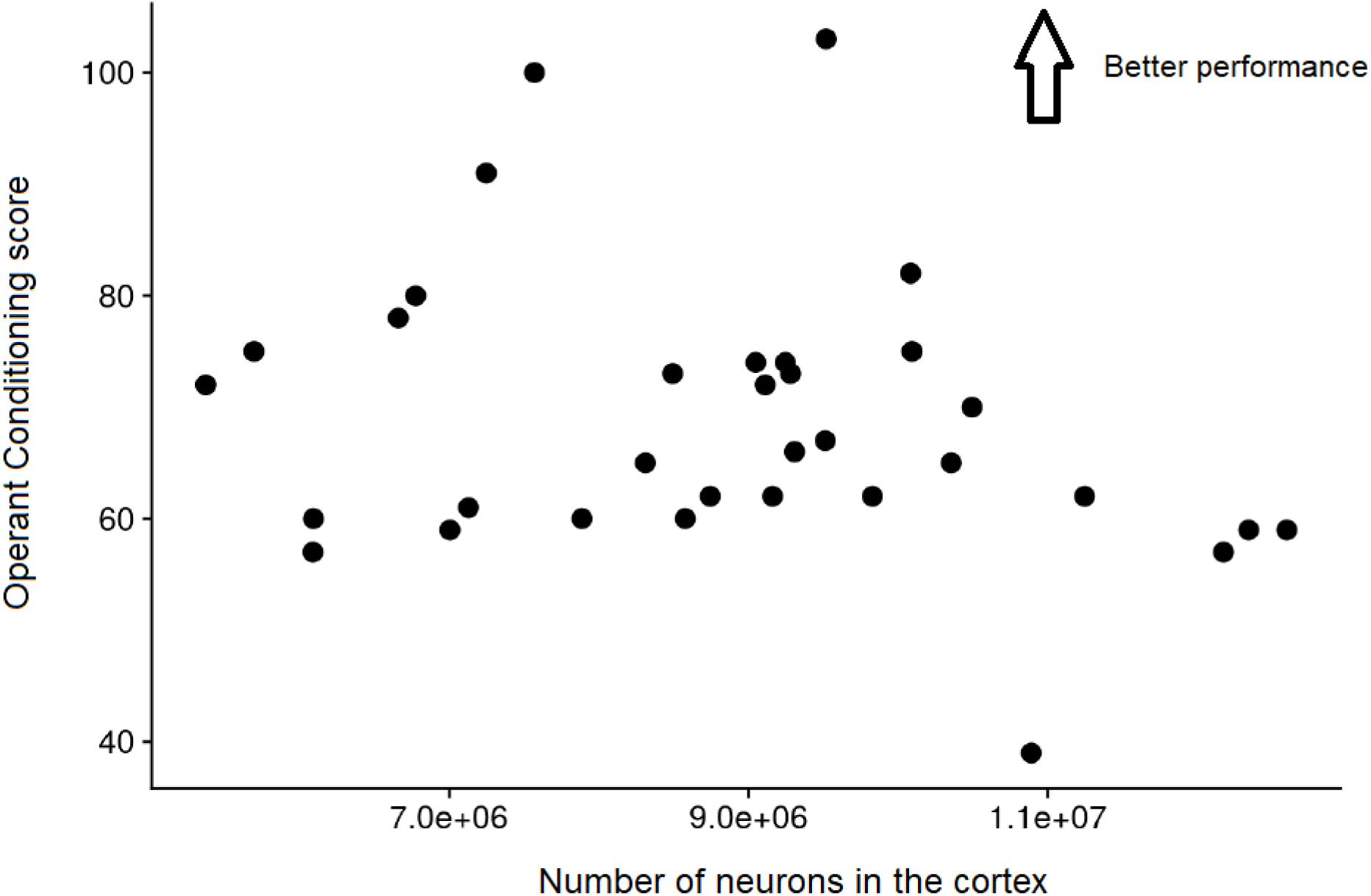
Lack of correlation between performance in the operant training and number of neurons in the cerebral cortex. (N = 32, ρ = −0.21, p = 0.15).

**Table 2.** Correlations between behavioral performance in the tasks and number of neurons in different structures. Each cell in the table shows the Spearman correlation and 95% confidence interval between behavioral performance and number of neurons for the various tasks and brain regions. No correlations but one, which was not an *a priori hypothesis* (Cerebral cortex, posterior x Rotarod) were significant at the 5% level. Sample size is 26–32, depending on the comparison.

|  |  | Number of Neurons |  |  |  |  |  |
| --- | --- | --- | --- | --- | --- | --- | --- |
|  | Cerebellum | Cerebral Cortex, anterior | Cerebral Cortex, posterior | Cerebral cortex, total | Hippocampus | Olfactory Bulb | Rest of Brain |
| **Rotarod** | 0.18<br>[-0.18, 0.49] | 0.3<br>[-0.05, 0.58] | 0.35<br>[0.01, 0.62] | 0.33<br>[-0.01, 0.61] | -0.09<br>[-0.42, 0.26] | 0.03<br>[-0.32, 0.37] | -0.16<br>[-0.48, 0.19] |
| **Puzzle Box** | 0.05<br>[-0.3, 0.38] | -0.2<br>[-0.51, 0.15] | -0.16<br>[-0.47, 0.2] | -0.17<br>[-0.49, 0.18] | -0.11<br>[-0.43, 0.24] | -0.13<br>[-0.45, 0.22] | 0.00<br>[-0.34, 0.34] |
| **Operant Conditioning AUC** | 0.16<br>[-0.19, 0.48] | -0.05<br>[-0.39, 0.3] | -0.13<br>[-0.45, 0.23] | -0.21<br>[-0.52, 0.14] | -0.14<br>[-0.46, 0.21] | 0.06<br>[-0.29, 0.4] | -0.19<br>[-0.5, 0.16] |
| **Morris Water Maze** | 0.17<br>[-0.18, 0.49] | 0.03<br>[-0.32, 0.37] | -0.32<br>[-0.6, 0.03] | -0.29<br>[-0.58, 0.06] | 0.03<br>[-0.32, 0.37] | -0.03<br>[-0.37, 0.32] | 0.03<br>[-0.32, 0.37] |
| **Hidden Peanut Butter Test** | -0.27<br>[-0.56, 0.08] | -0.26<br>[-0.55, 0.09] | -0.23<br>[-0.53, 0.13] | -0.19<br>[-0.5, 0.16] | 0.06<br>[-0.29, 0.39] | -0.13<br>[-0.45, 0.22] | -0.24<br>[-0.54, 0.12] |

To rule out the possibility that exposing mice to our battery of cognitive tests could alter the numbers of neurons, we compared the number of neurons of control and tested mice in two brain regions more likely to be affected by manipulations resulting in cognitive enrichment, namely the cerebral cortex and the hippocampus. Importantly, we did not find significant differences between control and tested animals in the number of neurons in the cortex **(Figure 12A)** and in the hippocampus **(Figure 12B)**.

**Figure 12:**
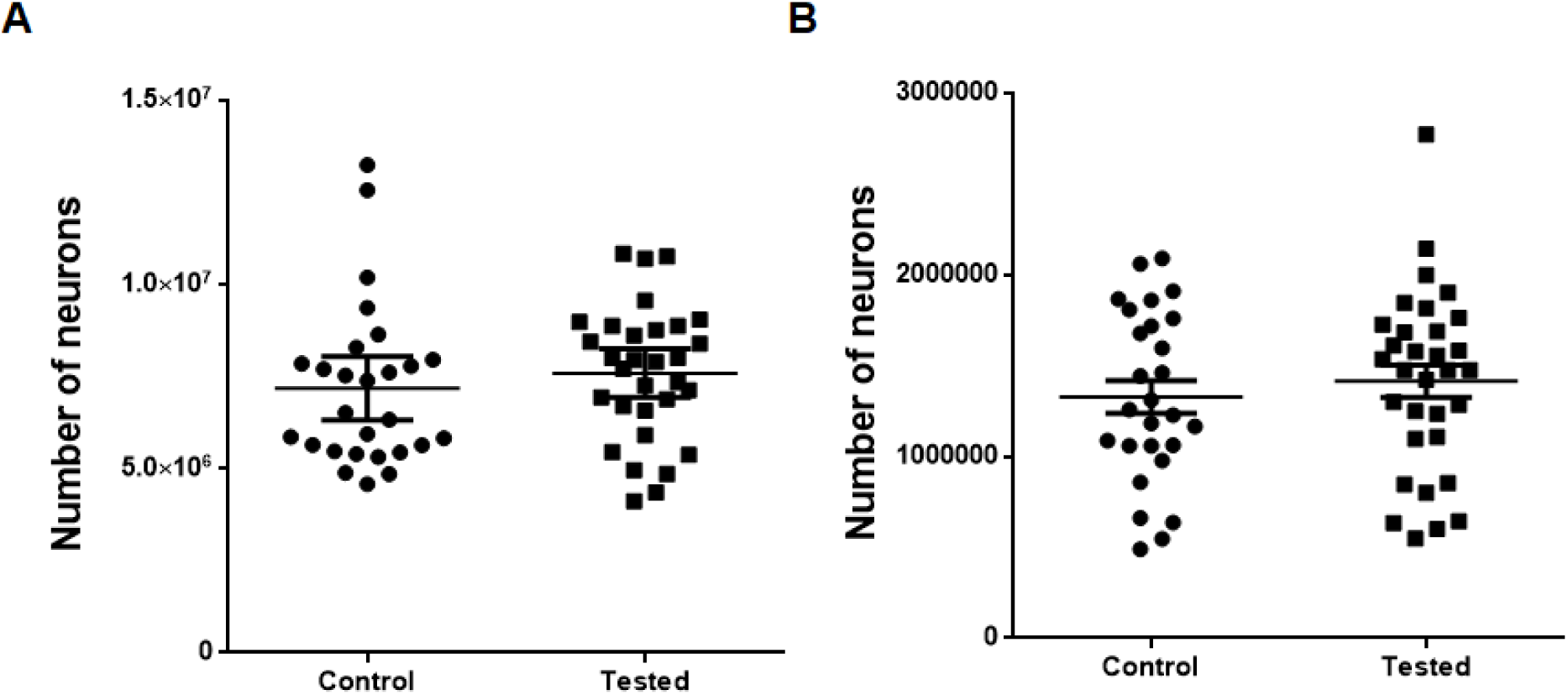
Testing mice through our cognitive battery does not change the number of neurons. Control and tested mice have similar number of neurons in the **(A)** cortex (n = 27 control and n = 31 tested, p = 0.43) and **(B)** in the hippocampus (n = 27 control and n = 32 tested, p = 0.49).

## Discussion

Although it is established that neuronal death is accompanied by worse behavioral performance across many tasks (Walker and Tesco, 2013; Kalaria et al., 2016), to our knowledge, our study is the first that attempts to investigate whether the number of neurons in normally-developed mice is associated with individual variation in behavioral performance. Our results suggest that naturally occurring variation in neuron number is not associated with variation in performance at the level of inexperienced individual animals within a species.

It could be argued that exposing mice to our cognitive battery changed neuronal numbers in ways that masked initial individual differences associated with behavioral performance. Indeed, many cognitive enrichment protocols have shown increased neurogenesis (Kempermann et al., 1998) and increased volume in specific brain regions (Maguire et al., 2000). However, it is unlikely that this played a role in our results, because going through our battery of behavioral tests did not influence neuronal number in the cortex **(Figure 12A)** nor in the hippocampus **(Figure 12B)** compared with untrained mice.

While we focused on neurons, it is known that other cell types, such as astrocytes and oligodendrocytes, also influence behavior. Han and colleagues (2013) investigated the effect of astrocytes on cognition, using a chimeric mouse model with grafted human astrocytes. They report that the chimeric mice perform better than control mice in an object location memory test and fear conditioning tasks. Moreover, myelination has been proposed as a correlate of intelligence (Miller, 1994). In any case, we did not find any correlation between individual behavioral performance and numbers of non-neuronal cells **(Table 3)**.

**Table 3.** Correlations between behavioral performance in the tasks and number of other cells (non-neuronal) in different structures. Each cell in the table shows the Spearman correlation and 95% confidence interval between behavioral performance and number of neurons for the various tasks and brain regions. No correlations but two (Morris Water Maze x Cerebellum and Morris Water Maze x Rest of Brain) were significant at the 5% level. Sample size is 26–32, depending on the comparison.

|  |  | Number of Other Cells |  |  |  |  |  |
| --- | --- | --- | --- | --- | --- | --- | --- |
|  | Cerebellum | Cerebral Cortex, anterior | Cerebral Cortex, posterior | Cerebral cortex, total | Hippocampus | Olfactory Bulb | Rest of Brain |
| **Rotarod** | 0.27<br>[-0.08, 0.56] | 0.23<br>[-0.13, 0.53] | 0.24<br>[-0.11, 0.54] | 0.33<br>[-0.01, 0.61] | 0.02<br>[-0.32, 0.36] | 0.11<br>[-0.24, 0.43] | -0.17<br>[-0.48, 0.19] |
| **Puzzle Box** | 0.12<br>[-0.24, 0.44] | -0.22<br>[-0.52, 0.13] | -0.06<br>[-0.39, 0.29] | -0.21<br>[-0.52, 0.15] | -0.11<br>[-0.44, 0.24] | -0.06<br>[-0.39, 0.58] | 0.23<br>[-0.12, 0.53] |
| **Operant Conditioning AUC** | 0.09<br>[-0.27, 0.42] | -0.02<br>[-0.36, 0.33] | 0.02<br>[-0.32, 0.36] | 0.06<br>[-0.28, 0.40] | 0.00<br>[-0.34, 0.35] | 0.30<br>[-0.05, 0.58] | -0.13<br>[-0.45, 0.22] |
| **Morris Water Maze** | -0.47<br>[-0.7, -0.15] | 0.00<br>[-0.34, 0.35] | -0.16<br>[-0.48, 0.19] | -0.14<br>[-0.46, 0.22] | 0.07<br>[-0.28, 0.41] | -0.10<br>[-0.43, 0.26] | 0.36<br>[-0.63, -0.02] |
| **Hidden Peanut Butter Test** | -0.20<br>[-0.51, 0.16] | -0.13<br>[-0.46, 0.22] | -0.16<br>[-0.48, 0.19] | -0.17<br>[-0.49, 0.18] | 0.05<br>[-0.30, 0.38] | -0.01<br>[-0.36, 0.33] | -0.11<br>[-0.44, 0.24] |

It remains possible that the number of neurons does correlate with behavioral performance, albeit weakly, and we did not find any significant correlation because of lack of statistical power. The confidence interval for some correlations include somewhat high values (e.g. number of neurons in the cerebellum and performance on the rotarod task shows a correlation of 0.18, with a 95% confidence interval going from −0.18 to 0.49). Conceivably, many factors are likely to influence the individual performance in a given task, such as stress reactivity and motivation to complete the task, multiple factors may compound where each one explains a small degree of the variance observed. However, the power for the sample size in our study is 80% for a linear correlation of about 0.5; for the sake of comparison, the statistical power is 35% for a correlation as weak as the one found for human brain volume and IQ in McDaniel, 2005 (a correlation of 0.29). In our study, however, we expected a correlation larger than that, for two main reasons: (1) as we have argued before, neuron number is more strongly correlated with behavioral performance across species than brain volume and (2) greater variability of subjects in human studies should add noise to the estimate, compared to the present study with lab mice.

Precise measures are essential to detect *small* effects. Regarding the behavioral measures, it is known that typically uncontrolled variables can have effects in animal behavior (Mandillo et al., 2008). The other indicator of interest, number of neurons, was measured using the isotropic fractionator, which relies on counting nuclei - stained with DAPI for total cells and with anti-NeuN for neurons - under a microscope (see Methods). The isotropic fractionator (see (Herculano-Houzel et al., 2015a)) is precise enough for detecting differences in numbers of brain neurons between closely related species that differ in behavioral performance: for instance, rats, which have just over twice as many cortical neurons as mice, learn operant conditioning tasks faster than mice (Jaramillo and Zador, 2014). The method also works for detecting differences in variance - isogenic C57BL/6 mice show less variation in neuron number between individuals than non-isogenic Swiss mice (unpublished observations). Nevertheless, the method has intrinsic error from the counting procedure and from sampling for immunocytochemical staining that makes estimates of neuron number vary around 10%-15% (unpublished results). Added together, the methodological variation could make small differences undetectable.

One could say, though, that whatever correlation might exist between the quantities analyzed here is likely to be small. Whereas the number of neurons is a strong predictor of cognitive performance across species (Harrigan and Commons, 2014; Dicke and Roth, 2016; Herculano-Houzel, 2017), our study suggests that this is not the case within a species. It may be simply that the magnitude of the differences within a species is not large enough to have a consistent, detectable impact on cognitive abilities.

A small or nonexistent correlation between behavioral performance and neuron number at the individual level invites a number of other possible explanations. First, we might not have tested the abilities for which having more neurons is beneficial. One recent study reviewed the literature in search of measures of cognitive performance of three animals - pigeons, corvids, and apes, as a function of increasing neuron number - on different tasks (Güntürkün et al., 2017). They found that the maximum level of performance is the same for all three animals in some of the tasks (e.g. short-term memory and abstract numerical competence). Remarkably, while pigeons do reach the same level of performance as primates and corvids on some tasks, they are slower to learn and have difficulty generalizing and transferring the associations to new contexts (Güntürkün et al., 2017). This suggests that the number of neurons might matter for the speed of learning, storage or generalization, but not to the final level of performance. Although the measures extracted from the cognitive tasks used in this study do take into account the learning rate, they also depend on the final level of performance.

One possibility is that having more neurons makes a larger difference for sensorimotor abilities, but not higher-order ones: more neurons could result in more precision in stimuli representation. Having more neurons available would decrease interference and overlap between stored patterns, reducing confusion in retrieval. Some studies lend support to this idea. A training-induced increase in the number of auditory neurons that respond preferentially to a given sound frequency correlates positively with a rat’s ability to identify said sound frequency (Polley et al., 2006). A similar relationship is observed between the primary motor area and motor skill in the corresponding areas of the body (Nudo et al., 1996). Remarkably, a manipulation that increases the number of neurons in the primary visual cortex of mice improves their visual discrimination (Fang and Yuste, 2017). However, we could not determine the number of neurons in sensorimotor cortices because their dissection is not reliable.

Another possibility is that what accounts for variation in behavioral performance is not variation of the number of neurons, but of their subcomponents - such as numbers of synapses, dendrites or channels - that determine information processing capacity. These components, in contrast to the number of neurons, are supposedly more plastic across the lifetime of an individual. It is likely that the way the brain is molded by experience is at least as important as more fixed biological/neuroanatomical parameters such as number of neurons for determining the cognitive skills of an individual. It is also possible that subcellular elements such as synapses and channels could process information by themselves, for example, by storing below-threshold voltage information (Sarpeshkar, 1998; Debanne et al., 2013) or a record of past firing activity in the channel’s state (Forrest, 2014).

Whatever the role of synaptic components in behavioral performance, neurons are a limiting resource for information processing. Dendrites, synapses and sodium-potassium pumps are all parts of neurons and theoretical predictions from information theory and biophysics suggest that is not possible to make these components more efficient: neural information processing already happens close to the minimum cost allowed by thermodynamics (Sterling & Laughlin, 2015). To the extent that a brain could have larger numbers of dendrites, synapses and pumps, those numbers would be limited by how many neurons the brain has.

In contrast, it is unlikely that numbers of glial cells are limiting to behavioral performance. These cells are only added in significant numbers, and very stable densities, to nervous tissue late in development, filling the neuronal parenchyma, which may account for the universality of the relationship between the glia/neuron ratio and neuronal density across mammals (Herculano-Houzel et al., 2014; Herculano-Houzel and Dos Santos, 2018). If that is the case, the number of glial cells depends on the number and size of neurons. Similarly, numbers of synapses might be an emergent property of the distribution of neurons and glial cells in the tissue, rather than under strong genetic control. The limited evidence available on synaptic densities across species suggests that they are fairly constant (Cragg, 1967; Schüz and Palm, 1989). In summary, most of the potential variation in the brain’s processing capacity across species and individuals comes from variation in the number and density of neurons. This contrasts with our results, which indicate that if a correlation does exist between neuronal number and individual cognitive capabilities in normally developed mice, then that the correlation is weak, at best.

In humans, intelligence is highly heritable (Plomin and von Stumm, 2018) and there is evidence that intraspecific brain size variation is heritable (Logan et al., 2016). Artificial selection experiments where changes in brain size are obtained in relatively few generations have successfully been conducted with rodents and fish (Kolb et al., 2013; Kotrschal et al., 2015). It would be informative to perform artificial selection studies for high or low cognitive performance in behavioral tasks and determine whether numbers of neurons in the corresponding brain structures respectively increased or decreased across generations.

Our finding of no obvious correlation between number of neurons and individual behavioral performance across many tasks in naive mice suggest that artificial selection for behavioral performance should not modify the average numbers of neurons that compose the brain of individuals in a population over time. This poses an interesting paradox for understanding the driving forces behind the evolution of larger brain size, for the lack of correlation undermines the customary prediction that brains with more neurons are adaptive and advantageous due to the presumably ensuing cognitive advantage that they bring about, and thus there should be positive selection pressure for larger numbers of neurons. If it is indeed the case that larger numbers of neurons are not predictive of any intrinsic cognitive individual advantage, then explaining mammalian brain evolution will require a major paradigm change in which the main driving force behind brain expansion is not cognitive performance directly, but possibly other factors such as decreased sleep requirement and a resulting alleviation in feeding pressure (Herculano-Houzel et al., 2015b). Future studies should aim to identify good predictors of variation in behavioral performance across individuals of the same species, such as genetic differences, numbers of synapses, numbers of specific glial cell subtypes, degree of myelination or dendritic arborization. We expect that, at the individual level, these variables may better predict the experience-dependent plasticity that underlies learning according to the individual history of interaction with the environment. Given that the brain is such a plastic organ, it seems plausible that activity-dependent self-organization could act as a stabilizing mechanism that buffers the brain against biological insults and other sources of variation that are likely to affect the number of brain neurons with which an individual is born.

## List of abbreviations

g: general intelligence factor
NeuN: neuronal nuclear antigen

